# Computational modeling of deep tissue heating by an automatic thermal massage bed: predicting the effects on circulation

**DOI:** 10.1101/2022.04.21.488942

**Authors:** Jacek P. Dmochowski, Niranjan Khadka, Luis Cardoso, Edson Meneses, Youngsoo Jin, Marom Bikson

## Abstract

Automatic thermal and mechanical massage beds support self-managed treatment, including reduction of pain and stress, enhanced circulation, and improved mobility. As the devices become more sophisticated (increasing the degrees of freedom), it is essential to identify the settings that best target the desired tissue. To that end, we developed an MRI-derived model of the lower back and simulated the physiological effects of a commercial thermal-mechanical massage bed. Here we specifically estimated the tissue temperature and increased circulation under steady-state conditions for typical thermal actuator settings (i.e., 45-65°C). Energy transfer across nine tissues was simulated with finite element modeling (FEM) and the resulting heating was coupled to blood flow with an empirically-guided model of temperature-dependent circulation. Our findings indicate that thermal massage increases tissue temperature by 3-8°C and 1-3°C at depths of 2 and 3 cm, respectively. Importantly, due to the rapid (non-linear) increase of circulation with local temperature, this is expected to increase blood flow four-fold (4x) at depths occupied by deep tissue and muscle. These predictions are consistent with prior clinical observations of therapeutic benefits derived from spinal thermal massage.

## Introduction

Direct application of heat to the skin is known to increase blood flow (1, 2), an essential physiological response that also removes excess heat actively produced during metabolism and exercise. The increased circulation produced by passive heating has been exploited in multiple therapeutic settings, most notably in the management of pain (3). Medical devices that provide thermal stimulation are becoming increasingly sophisticated. For example, automatic thermal-mechanical massage beds are equipped with multiaxis traction and far-infrared thermal projectors (4–6). Such devices allow in-home, self-managed treatment that may potentially leverage the numerous degrees of freedom governing the stimulation “dose”. In theory, stimulation parameters such as temporal dynamics (pulsed versus continuous), intensity (temperature), duration, and position may be individualized to maximize therapeutic benefit. Understanding the relationship between these numerous parameters and the resulting physiological changes is critical to such optimization efforts.

A central question in thermal therapies pertains to target engagement: given a targeted region (e.g. a specified muscle group), what is the required temperature that must be applied to the surface in order to produce the desired increase in blood flow to the target? Addressing this question requires knowledge of the temperature distribution produced throughout the volume during passive heating, as well as the quantitative relationship between the temperature distribution and resulting change in circulation. Recent work indicates that the increase in circulation during passive heating is largely driven by *local* temperature gradients (7), implying that the magnitude of circulation increase may be predicted with knowledge of the local temperature. To estimate the temperature distribution produced by a given thermal device, computational modeling that accounts for the heterogeneous anatomical structures of the treated area may be leveraged. Indeed, computer-based simulations are standard tools across medical device design (such as neuromodulation) and increasingly rely on detailed image (e.g., MRI) derived representation of the underlying anatomy (8–11).

Here we performed a realistic simulation of tissue heating and associated circulation changes produced by a commercial thermo-mechanical massage bed. We optimized MRI imaging to resolve key structures of the back, developed a 3D model of the relevant tissues, modeled the action of thermal actuators under static conditions, and employed finite element modeling to predict the intensity and distribution of temperature in the lumbar region. We then leveraged prior work relating local temperature to circulation to construct a model of blood flow changes during thermal massage. Our findings indicate that temperature increases up to 3°C are achievable at deep (3 cm) regions of the lumbar back with conventional actuator settings. Moreover, our findings suggest that blood flow can be increased four-fold at depths of 2-3 cm, corresponding to the lumbar musculature. These findings are consistent with clinical observations performed with the same thermo-mechanical massage bed (4–6, 12).

## Methods

### MRI scanning and tissue segmentation

We performed T2-weighted lumbar spine magnetic resonance imaging (MRI) of a healthy male adult (BMI: 25 kg/m^2^; age: 42 years) with a 3T Siemens MAGNETOM Prisma scanner equipped with a CP Spine array coil (Siemens Healthineers, PA, USA). A sagittal slice is shown in Fig. 1a. The parameters of image acquisition were set according to: TE = 99 ms, TR = 7040 ms, flip angle = 130°, FOV = 256 mm, SNR = 1, in-plane resolution = 256 x 256, slice thickness = 1 mm; and voxel size: 1 x 1 x 1 mm.

**Fig. 1.**
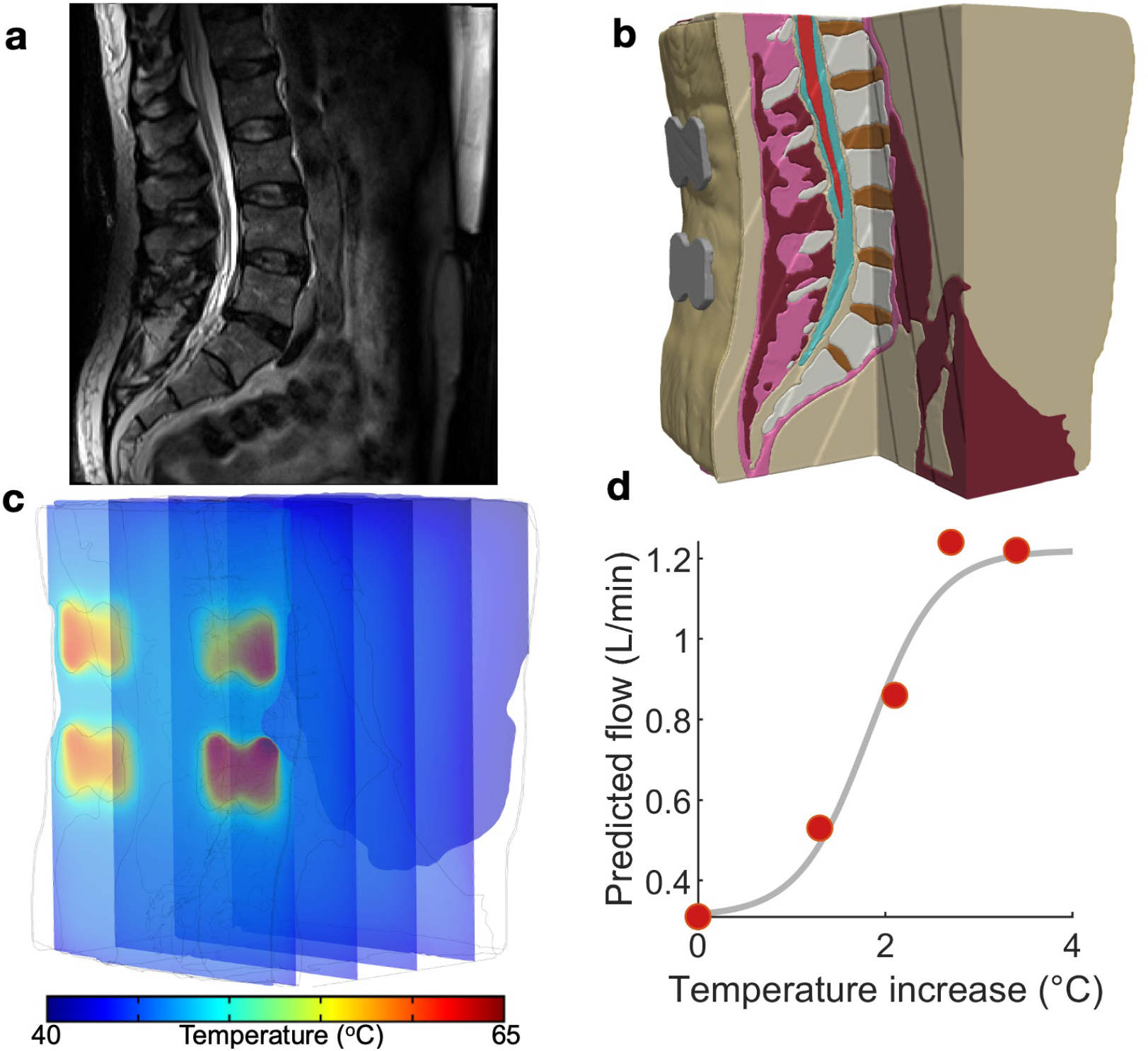
Constructing a computational model to predict tissue temperature and blood flow during thermal massage. **(a)** Sagittal slice of a T2-weighted anatomical MRI used to construct the computational model developed in this study. **(b)** Based on image contrast, the volume was segmented into nine tissues that were then endowed with physical properties governing the physiological response to an external heat source. **(c)** The Pennes bio-heat equation was solved with a finite element model (FEM) solver, yielding the temperature during thermal massage throughout the volume. The locations of the four actuators are apparent from the temperature “hot-spots” in the superficial slice. **(d)** In order to translate model derived tissue temperature to the predicted changes in blood flow, we constructed an empirically-guided sigmoidal model that outputs the expected blood flow based on *in situ* tissue temperature.

Based on image intensity, the MRI was segmented into nine tissue masks: skin, subcutaneous fat, soft tissue, muscles, intervertebral disc, vertebrae, epidural fat, cerebrospinal fluid, and spinal cord. Manual tissue segmentation was carried out by first thresholding the images, followed by morphological filtering such as flood fill, dilation, and erosion, performed in Simpleware ScanIP (Synopsys Inc., CA, USA). To ensure tissue continuity and improve segmentation accuracy, the data was extensively visualized and compared with the anatomy of the subject’s spine while performing manual adjustments to the tissue masks. Based on the results of segmentation, the thicknesses of the tissues comprising the imaged region were measured as: (skin) 1.1 mm, (subcutaneous fat) 13 mm, (muscle) 42 mm, (soft tissue) 2.8 mm, (epidural fat) 2.2 mm, (CSF) 2.9 mm, and (spinal cord) 1.9 mm. Thus, the muscle (i.e., the presumed target of thermal massage) occupied a region approximately 1.5 - 5.5 cm from the surface.

### Finite element modeling

The tissue masks generated by the segmentation process were utilized to construct a finite element model (FEM) of the lumbar back. To that end, we employed the Simpleware Scan IP software running the tetrahedral voxel-based meshing algorithm to generate a FEM consisting of more than 3.38 million tetrahedral elements. The resulting model was then integrated with separate domains that captured the position, orientation, and geometry of a thermal massage device. In particular, we modeled the characteristics of the CGM MB-1901 massage bed (CERAGEM Co. Ltd., Cheonan, Korea), which is equipped with two types of heat sources: (i) a contoured heating “mat” that resides directly under the bed surface and provides a diffuse layer of heat, and (ii) four actuators (“rollers”) that make direct contact with the body to provide more punctate and intense thermal stimulation. The device contains four actuators arranged as the vertices of a rectangle: the lateral distance between actuators is 56 mm, and the vertical distance between top and bottom actuators is 32 mm. Specifically, we modeled cross-sections of the four actuators (width: 65 mm, diameter of the circular ends: 45 mm, spacing between the two circular ends: 30 mm) in Solidworks (Dassault Systems, MA, USA) and imported the components into Simpleware ScanIP for positioning and meshing. The resulting model was then imported into COMSOL Multiphysics 5.5 (COMSOL Inc., MA, USA) to solve the Pennes’ bio-heat transfer equation

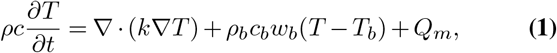

where *ρ* is the tissue density (units of kg/m^3^), *c* is the specific heat capacity (J kg^−1^ K^−1^), *T* is the temperature (K), *t* denotes time (s), *k* is the thermal conductivity (W m^−1^ K^−1^), *ρ_b_* is the density of blood (kg m^−3^), *c_b_* is the specific heat capacity of blood (J kg^−1^ K^−1^), *w_b_* is the perfusion rate of blood (1/s), *T_b_* is blood temperature (K), and *Q_m_* is the rate of metabolic heat generation (W/m^3^). The solution was carried out under steady state conditions.

The physical and thermal properties of the model tissues were assigned according to the values listed in Table 1, and were obtained from a survey of prior literature (13–19). In order to solve the bio-heat equation (1), the temperature at the model’s external boundaries was fixed to core body temperature (37°C). We assumed the absence of convective gradients across all internal tissue boundaries. Convective heat loss from model boundaries to the environment was modeled as:

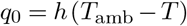

where *q*_0_ (units of W m^−2^) is the convective heat flux, *h* = 5 W m^−2^ K is the heat transfer coefficient for the tissues comprising the model, and *T*_amb_ is the ambient temperature (i.e., 25 °C). The initial temperature of the tissues was set to 37°C. The temperature of the actuators was fixed within a simulation to a value ranging from 45 to 65 °C. The external boundary of the skin making contact with the bed surface, corresponding to the heating mat, was assigned a temperature of 40°C.

**Table 1.** Thermal properties of the tissues and sources comprising the computational model developed here to estimate temperature gradients during thermal massage.

| Material<br>units | $k$<br>$\text{Wm}^{-1}\text{K}^{-1}$ | $\rho$<br>$\text{kg m}^{-3}$ | $c$<br>$\text{Jkg}^{-1}\text{K}^{-1}$ | $\rho_b$<br>$\text{kg m}^{-3}$ | $c_b$<br>$\text{Jkg}^{-1}\text{K}^{-1}$ | $w_b$<br>1/s | $T_b$<br>K | $Q_m$<br>$\text{W/m}^3$ |
| --- | --- | --- | --- | --- | --- | --- | --- | --- |
| skin | 0.37 | 1100 | 3391 | 1057 | 3600 | 0.0004 | 309.7 | 457 |
| muscle | 0.47 | 1142 | 3432 | 1057 | 3600 | 0.0004 | 309.7 | 457 |
| soft tissue | 0.47 | 1142 | 3432 | 1057 | 3600 | 0.0004 | 309.7 | 457 |
| vertebrae | 0.32 | 1908 | 1313 | 1057 | 3600 | 0.0003 | 309.7 | 342 |
| i.v. disc | 0.49 | 1100 | 3568 | 0 | 0 | 0 | 309.7 | 0 |
| subcut.<br>epidural<br>fat | 0.21 | 1142 | 2348 | 1057 | 3600 | 0.00008 | 309.7 | 302 |
| CSF | 0.57 | 1007 | 4096 | 0 | 0 | 0 | 310 | 0 |
| spinal cord | 0.51 | 1075 | 3630 | 1057 | 3600 | 0.008 | 309.7 | 9121 |
| actuators | 237 | 2710 | 900 | - | - | - | - | - |

### Model of temperature-dependent circulation

In order to estimate the magnitude and distribution of blood flow produced during thermal massage, we developed a non-linear model for mapping the *in situ* tissue temperature to the resulting blood flow. The model was guided by simultaneous empirical measurements of temperature and blood flow in the human leg during heat stress as conducted by Chiesa and colleagues (7). The general form of the model developed here was given by:

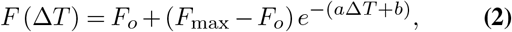

where *F* is the blood flow (L/min), Δ*T* is the change in temperature at the site of the vessel (°C), *F_o_* is the blood flow in the absence of exogenous heating, *F*_max_ is the maximum achievable blood flow due to the physical limitations of the cardiovascular system, and the free model parameters *a* and *b* were estimated by fitting the model to the empirical measurements reported by Chiesa and colleagues. Namely, the following (*T,F*) pairs of measurements were reported, where *T* was taken here to be the temperature in the muscle of the leg (i.e., the deepest measurement taken and the one closest to the site of the vessel) and *F* was taken to be the empirical blood flow in the central femoral artery (CFA): (34.9°C, 0.31 L/min), (36.2°C, 0.53 L/min), (37.0°C, 0.86 L/min), (37.6°C, 1.24 L/min), and (38.3°C, 1.22 L/min).

From these empirical measurements, the values *F_o_* = 0.31 L/min and *F*_max_ = 1.22 L/min were assigned to the baseline and maximum blood flow, respectively. We then performed conventional least-squares fitting employing the Matlab (Mathworks, MA, USA) function *lsqcurvefit*. The minimization procedure yielded values of *a* = −2.67 and *b* = 4.88, producing excellent fits to the empirical measurements while capturing a sigmoidal transition from baseline to maximal flow (Fig. 1d). The resulting model was then employed to transform local temperature to the corresponding blood flow.

## Results

We simulated the action of a commercial automatic thermal massage device, where contact heating was delivered through (i) a contoured plane (“mat”) at 40°C covering the full extent of the lower back, and (ii) four actuators (“rollers”) with a diameter of 4.5 cm and a height of 1 cm. The actuators were arranged on the back as the vertices of a rectangle (Fig. 1b,c). The actuator temperature was varied from 45 to 65 °C in increments of 5°C. For each intensity, the temperature distribution throughout the lumbar region was numerically computed. The resulting solution was then coupled to the predicted change in blood flow. Our objective was to determine the magnitude of the circulation increase as a function of tissue depth and actuator temperature.

### Spatial distribution of lower back temperature during thermal massage

The spatial distribution of temperature at a sagittal slice immediately anterior to a pair of actuators is shown in Fig. 2a (shown here for an actuator temperature of 65°C), while the contours of the temperature field are depicted in Fig. 2b. The spatial dynamics of the temperature increase are apparent, and it is clear that the largest changes occur near the stimulation site (note the higher density of contour lines). To quantify the effect of tissue depth on achieved temperature, we analyzed temperature immediately anterior to the centroid of an actuator, as a function of depth. The tissue temperature exhibited an exponential decay with depth (Fig. 2e, colors indicate the input temperature, markers indicate depths of 1, 2, 3, and 4 cm). With an actuator temperature of 45 °C, the temperature at depths of 1, 2, and 3 cm into the tissue was 42.2 °C, 39.6 °C, and 38.0 °C, respectively (Fig. 2e, blue). With an increased roller temperature of 65°C, the corresponding temperatures were 54.9°C, 45.5°C, and 40.6°C (Fig. 2e, green). Thus, the temperature gradient at a depth of 2 cm ranged from 3-8°C, while the gradient at 3 cm was 1-3°C.

**Fig. 2.**
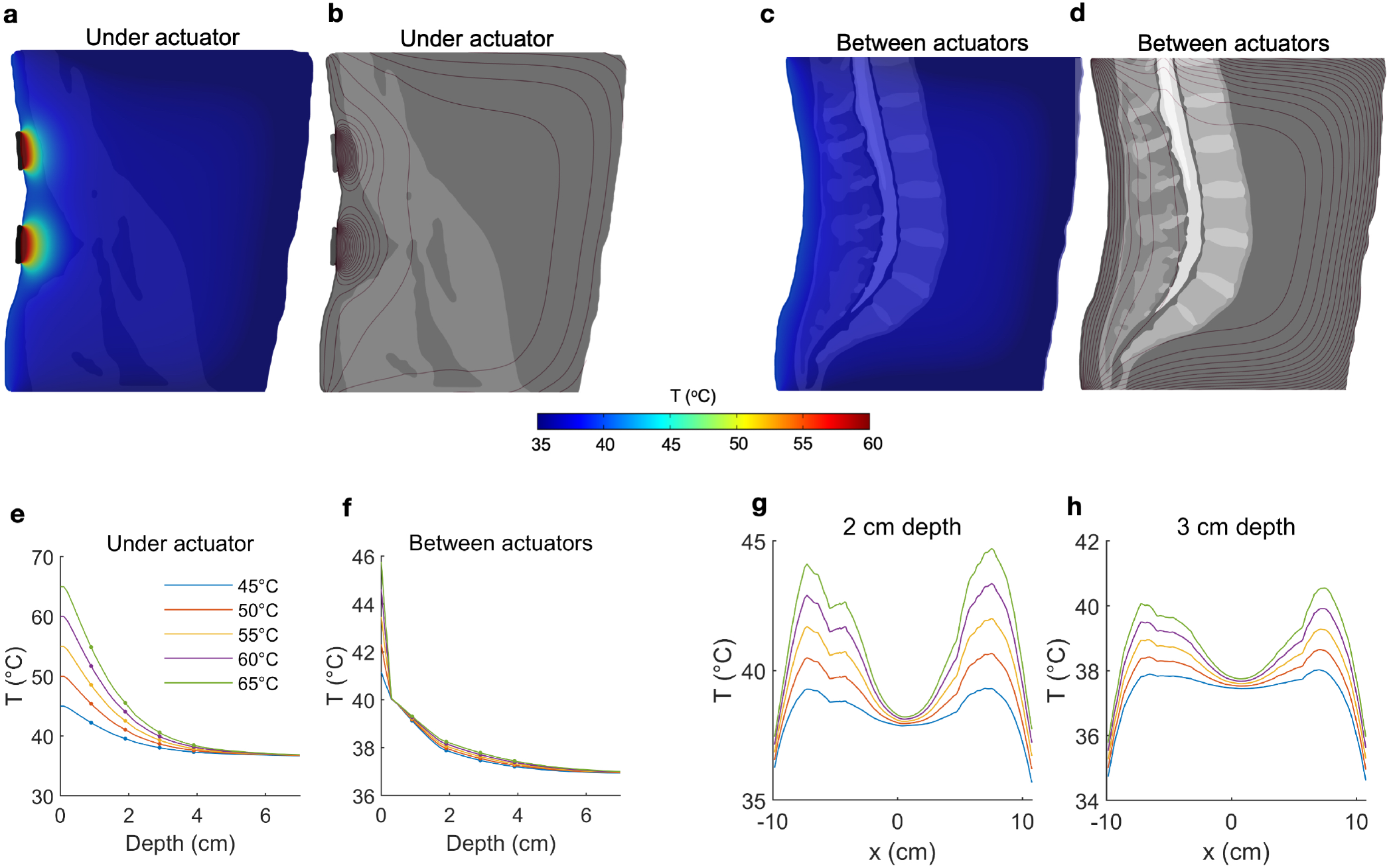
Quantifying temperature increases during thermal massage. **(a)** Temperature of a sagittal slice of the volume during thermal massage with the actuators set to 65°C. The selected slice was located directly anterior to the actuators. The resulting temperature decayed exponentially with distance from the heating elements. **(b)** The contours of the temperature distribution shown in (a). **(c)** Temperature of a sagittal slice located between actuators. The resulting temperature increase was markedly lower. **(d)** Contours corresponding to the temperature distribution in (c). **(e)** The temperature directly anterior to an actuator as a function of depth, shown for five different actuator temperatures. At a depth of 2 cm, temperature increased by 3-8°C. The temperature increase at 3 cm depth was between 1-3°C. **(f)** Between actuators, the achieved temperature increase was greatly dampened, with an increase of only 1 °C at a depth of 2 cm. **(g)** The tissue temperature distribution from left to right, shown for a fixed depth of 2 cm. The proximity of the tissue to the actuator placement determines the level of heating (actuators are positioned at ±6 cm). **(h)** Same as (g) but now for a depth of 3 cm, showing the increment in deep tissue temperature for different actuator settings.

Next we considered the effect of passive heating on tissue that is laterally displaced from the contact site. Fig. 2c depicts the spatial distribution at a sagittal slice positioned at the midline (i.e., between pairs of actuators, 6 cm from the actuator centroid and 3.75 cm from the most lateral edge). At an actuator temperature of 45°C, tissue temperatures were computed as 39.1°C, 37.9°C, and 37.5°C at depths of 1, 2, and 3 cm, respectively (Fig. 2f, blue). The corresponding temperatures with an actuator temperature of 65°C were only marginally higher: 39.3°C, 38.3°C, and 37.8°C (Fig. 2f, green), indicating that increasing source intensity does not compensate for a lateral displacement of the target.

To better characterize the effect of thermal massage on tissue temperature, we also analyzed the temperature across the full lateral extent of the back, but at fixed depths of 2 cm (Fig. 2g) and 3 cm (Fig. 2h). We considered a vertical location matching one (horizontal) pair of actuators. The resulting temperature profile confirmed the preferentially local action of the actuators, as the temperature fell to near mean body temperatures at the midline (37.9°C at a depth of 2 cm, and 37.5°C at a depth of 3 cm, Fig. 2g, h). With an actuator temperature of 45°C evaluated at a depth of 2 cm, the temperature fell from a peak of 39.3°C to 38.9 °C with a lateral displacement of 1 cm towards the midline, to 37.9 °C with a displacement of 2 cm, and to 36.4 °C with a displacement of 3 cm (Fig. 2g, solid blue). With an elevated actuator temperature of 65°C, the corresponding values were 44.6°C (peak), 43.8°C (1 cm displacement), 41.6°C (2 cm), and 38.3°C (3 cm; Fig. 2g, solid green). Evaluated at a depth of 3 cm, the peak temperature with actuators set to 45°C was 38.0°C, which fell to 37.7°C, 37.1°C, and 35.2°C when laterally displaced by 1, 2, and 3 cm, respectively. When simulating an actuator temperature of 65°C, lateral displacements of 1, 2, and 3 cm reduced a peak temperature of 40.5°C to 40.2°C, 39.2°C and 36.7°C, respectively (Fig. 2h).

### Predicted blood flow increases in the lower back during thermal massage

In order to predict the magnitude of increased blood flow during thermal massage, we developed a simple non-linear model of the relationship between *in situ* tissue temperature and blood flow. The model was based on concurrent empirical measurements of temperature and blood flow in the human leg during heat stress, where flow was found to non-linearly increase from a baseline value of 0.31 L/min to a maximum value of 1.22 L/min (Fig. 1d; see *Methods* for details of the empirical data and model fitting procedure). We employed the resulting sigmoidal model to transform the computationally derived tissue temperature values (shown in Fig. 2) to estimates of blood flow in the lower back during thermal massage.

The spatial distribution of predicted blood flow is shown in Fig. 3a,b for a sagittal slice immediately anterior to a (vertical) pair of actuators, and in Fig. 3c,d for a mid-sagittal slice (both shown for an actuator temperature of 65°C). Upon comparing Fig. 3a to Fig. 2a, it is evident that blood flow exhibits a more favorable decay with depth.

**Fig. 3.**
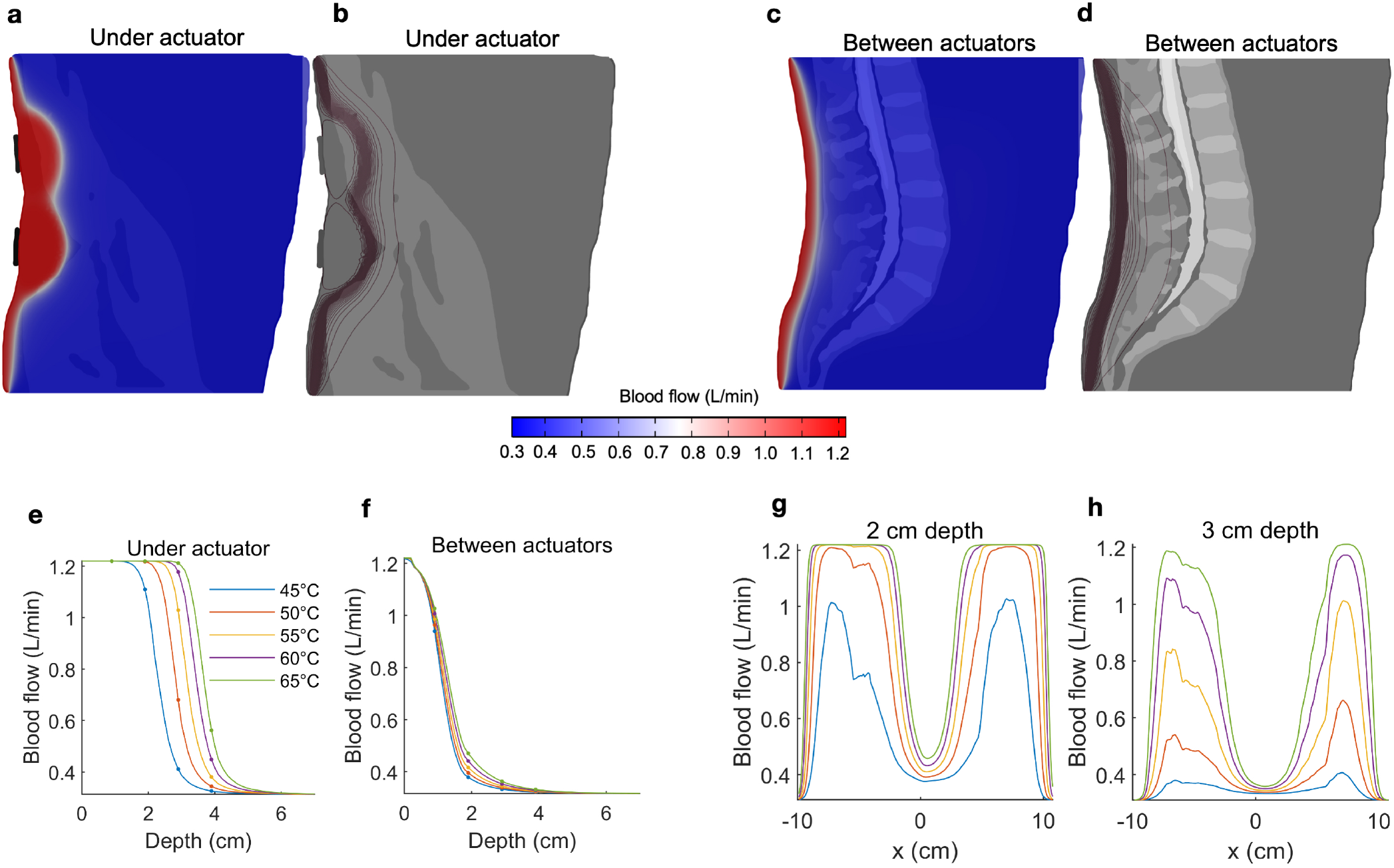
Predicting blood flow increases during thermal massage. **(a)** The predicted blood flow at a sagittal slice of the back, immediately anterior to the actuator. The actuator temperature was set to 65°C. Compared to temperature (Fig 2), pronounced increases in blood flow are observed at greater depths. **(b)** Contours of the blood flow distribution in (a), where the concentration of the contours indicates a sharp transition region from high to low blood flow. **(c)** The predicted blood flow at a sagittal slice located between actuators. The increase in blood flow is visibly dampened. **(d)** Contours of the blood flow distribution in (c), where the transition region is observed at a relatively shallow depth compared to (b). **(e)** The blood flow achieved immediately anterior to the actuators, shown for five different actuator intensities. Due to the sigmoidal nature of the temperature-circulation relationship, the falloff with depth occurs at a larger depth (between 2-4 cm, depending on actuator setting). Four-fold increases in circulation are predicted at depths of 2-3 cm. **(f)** Same as (e) but now shown for a sagittal slice located between the actuators. The increase in blood flow is no longer apparent beyond a depth of 3-4 cm. **(g)** The distribution of blood flow as a function of left-to-right position, shown for a fixed 2 cm depth. The predicted blood flow saturates (4x increase relative to baseline) across large sections of the back. **(h)** Same as (g) but now shown for a depth of 3 cm. When located directly anterior to the actuator, the predicted blood flow reaches a three-fold increase at an actuator temperature of 55°C.

At an input temperature of 45°C, the predicted blood flow was 1.22 L/min, 1.11 L/min, 0.41 L/min, and 0.33 L/min at depths of 1, 2, 3, and 4 cm, respectively (Fig. 3e, blue line). With an elevated input temperature of 65°C, the predicted flow saturated at depths of up to 3 cm (1.22 L/min), before sharply falling to 0.56 L/min at a depth of 4 cm (Fig. 3e, blue line). It is evident that the nonlinearity of the relationship between temperature and blood flow leads to abrupt transitions between low, baseline flow and large, saturated flow. Indeed, the sharp sigmoidal transition (Fig 1d) predicts that large increases in blood flow are achievable at depths of 3 cm, where a flow of 1.03 L/min is predicted with an actuator temperature of 55 °C (Fig. 3e, yellow line).

When considering blood flow as a function of lateral displacement from the center of the actuator, we observed that, given a sufficiently high input temperature (i.e., 55 °C), the blood flow at a depth of 2 cm saturated across a large lateral extent of the lower back (Fig. 3g). Even at a depth of 3 cm, the predicted flow produced by an actuator temperature of 55°C reached a three-fold increase at locations immediately anterior of the actuators (Fig. 3g). Overall, the effective sensitivity of blood flow to distance from the heat source is less than that observed with temperature.

## Discussion

We employed finite element modeling to estimate the physiological effects of an automatic thermal massage bed on the lumbar back region. The model predicts that it is feasible to produce large (e.g. four-fold) increases in circulation at relatively deep locations with high but practical actuator temperatures. The basis of this finding is that blood flow increases quickly and saturates at *in situ* temperature gradients of approximately 3° (7). Specifically, our FEM model predicts that such temperature increases are attainable at depths of 2 to 3 cm with actuator temperatures between 55° and 65°, values that overlap with those tested in earlier studies (5, 6). Thus, despite the exponential decay of temperature with depth (Figure 2), there is a sufficient gradient produced at the depths occupied by muscle tissue (1.5 - 5.5 cm) to modulate blood flow to the muscles of the lumbar region, which represent a common target of therapeutic massage beds.

Thermotherapy (3, 20) and massage (21, 22) have an extensive history in wellness and medicine but recent advances in automatic massage beds suggest new potency for self-managed treatment. Evidently, the therapeutic outcomes of thermal-mechanical massage will depend on the technical features of the mechanical and thermal actuators, as well as (individualized) anatomy and physiology. Notice, for example, that the position of tissues relative to the actuators had a strong influence on the resulting temperature and blood flow changes (Figs. 2 and 3). Knowledge of the distance and orientation of the target relative to the actuator(s), which will generally vary with the body composition of the individual, may be leveraged by future devices to personalize treatment. For example, the actuator temperature and treatment duration required to achieve a certain benchmark circulation increase may be determined automatically, potentially systematizing treatment.

Computational models of medical devices relate device features (exogenous and set by the operator) to the resulting physiological changes, which in turn govern therapeutic actions. Therefore, the effects on heating and circulation predicted here have direct implications for understanding (and further optimizing) results from clinical trials using the same device. As both temperature and blood flow are known to impact immune function, antioxidant and anti-inflammatory processes, and autonomic function (23–28), these predictions help to explain the clinical biomarkers observed during application of the device modeled here (4–6). In turn, these biomarkers provides a mechanistic substrate for thermal massage therapy in pain (4, 6).

As with any model, our predictions are subject to assumptions, including the transfer function employed here to relate temperature increases to blood flow. Valuable next steps will include (1) directly verifying model predictions by physiological measurements conducted either during or after thermal massage; and (2) designing and validating improvements in automatic massage bed operation. The present study assumed a static (stationary) regime when solving for the temperature during thermal massage. The rationale for this assumption is that the typical duration of thermal massage (e.g. tens of minutes) is expected to be larger than the time constants defining the transients of the bio-heat equation (29). In any case, the predictions made here correspond to the physiological effects produced when transients have passed and steady-state values are achieved. Nevertheless, the addition of a dynamic component to the simulation may allow one to identify potential advantages of pulsed stimulation, if for example, the vascular response is reinforced by a time-varying heat source marked by frequent onsets. Moreover, the mechanical deformations that occur in the lumbar region during thermal-mechanical massage may bring target tissues closer to the heat source during certain periods of the treatment, potentially increasing the improvement in circulation that was measured in a static configuration here.

Future modeling efforts should therefore integrate the mechanical effects of massage with the simultaneous temperature changes produced by thermotherapy in a multi-physics simulation that captures the potentially synergistic property of thermal massage. This may reveal that thermal stimulation modulates mechanical tissue properties and promotes the beneficial effects of tissue manipulation. Such symbiotic outcomes of thermal-mechanical massage are expected to translate to improved clinical outcomes, and will rely on computational modeling to design therapies that optimize the interplay between heating and mechanical forces.

## Author Contributions

JD designed the research, performed the research, and wrote the manuscript. NK designed the research, performed the research, and wrote the manuscript. LC designed the research. EM performed the research. YS designed the research. MB designed the research and wrote the manuscript.

